# Neural Network Poisson Models for Behavioural and Neural Spike Train Data

**DOI:** 10.1101/2020.07.13.201673

**Authors:** Moein Khajehnejad, Forough Habibollahi, Richard Nock, Ehsan Arabzadeh, Peter Dayan, Amir Dezfouli

**Author notes:** Authors contributed equally to this manuscript (Moein Khajehnejad, Forough Habibollahi).

## Abstract

One of the most important and challenging application areas for complex machine learning methods is to predict, characterize and model rich, multi-dimensional, neural data. Recent advances in neural recording techniques have made it possible to monitor the activities of a large number of neurons across different brain regions as animals perform behavioural tasks. This poses the critical challenge of establishing links between neural activity at a microscopic scale, which might for instance represent sensory input, and at a macroscopic scale, which then generates behaviour. Predominant modeling methods apply rather disjoint techniques to these scales; by contrast, we suggest an end-to-end model which exploits recent developments of flexible, but tractable, neural network point-process models to characterize dependencies between stimuli, actions, and neural data. We apply this model to a public dataset collected using Neuropixel probes in mice performing a visually-guided behavioural task as well as a synthetic dataset produced from a hierarchical network model with reciprocally connected sensory and integration circuits intended to characterize animal behaviour in a fixed-duration motion discrimination task. We show that our model outperforms previous approaches and contributes novel insights into the relationships between neural activities and behaviour.

## 1 Introduction

Recent developments in neural recording techniques such as Neuropixel probes allow the activities of large numbers of neurons across the brain to be monitored as animals perform behavioural tasks [Jun et al. 2017]. This allows us to study how the brain represents past and present sensory inputs across areas, how these representations evolve over time and ultimately lead to behaviour.

Very coarsely, supervised, reinforcement learning and unsupervised methods have been applied to examine the relationships between activity and behaviour [Paninski, Pillow, and Lewi 2007; Mante et al. 2013; Ganguli and Sompolinsky 2012; Kass, Eden, and Brown 2014; Sussillo 2014; Richards et al. 2019; Schaeffer et al. 2020]. Encoding and decoding models are examples of supervised learning. Encoding models use linear or non-linear methods to predict the activity of individual neurons based on task-related variables such as stimuli, actions, rewards and the like. Decoding models use linear or non-linear methods to predict the values of these task-related variables from the conjoint activities of multiple neurons within or between areas. Both types of model have been hugely influential from the earliest days of the application of computational methods to understand neural representation and processing [Dayan and Abbott 2001; Rieke et al. 1999; Kass, Eden, and Brown 2014; Meyer et al. 2017]. However, simple reflections of the computational constraints of the task are often insufficient to capture complex neural representations of the sensory inputs and actions that are distributed across different brain regions [Steinmetz et al. 2019] and can also evolve over time in ‘null’ neural modes that have no behavioural consequence. Moreover, neural responses are variable and a population’s response will often differ from trial to trial and over time, even under the same experimental conditions [Shadlen and Newsome 1998]. Most traditional approaches do not generalize to these more naturalistic conditions where trials with identical stimuli do not exhibit identical behavioural schemes [Goris, Movshon, and Simoncelli 2014; Churchland et al. 2010]. Hence, they can not account for the temporal irregularities in behaviour and neural recordings between trials.

Reinforcement learning (or in some cases, supervised learning) methods have more recently been used to learn potentially complex feedforward and recurrent neural network (RNN) models that are themselves capable of performing the same behavioural task as the subjects, based on assumptions about sensory noise and processing architecture [Barak 2017; Sussillo 2014; Richards et al. 2019; Schaeffer et al. 2020; Yamins et al. 2014]. These have been highly revealing about neural coding. However, they are also ill-suited to capture the myriad complexities of null modes, or the particular sub-optimalities expressed by individual subjects, reflecting their particular incompetence, training history and more.

Unsupervised methods have also been applied – often ways of mapping very high dimensional population activity into lower dimensional spaces [Paninski et al. 2010; Cunningham and Yu 2014; Whiteway and Butts 2019; Yu et al. 2006]; with the structure in these spaces, and perhaps the dynamical evolution of the states in these spaces, subsequently being related to task variables. These methods are typically useful since the number of dimensions of task input and/or output variability is often rather modest, implying that much of the high dimensional space that could potentially be occupied is either empty or at least not relevant for behaviour. However, they typically have to use intrinsic metrics such as variance to specify which low dimensional projections should be considered – and this again begs the question as to what is important.

Of course, there are methods that combine various of these approaches [Kobak et al. 2016; Kriegeskorte and Kievit 2013], and continual innovations. Along these lines, [Dezfouli et al. 2018] recently suggested a combined approach, with an RNN being trained to tie fMRI BOLD activity across the brain directly with ongoing behaviour. fMRI data allowed for a form of model inversion, pinning down the RNN state and so implying how behaviour would be realized neurally. However, this approach is licensed by the invertibility that is at least plausible because of the high dimensionality of fMRI. It is not currently guaranteed to be available on a trial-by-trial basis in neural recordings.

Here, motivated by previous approaches and recent neural network point-process models [Omi, Aihara, et al. 2019], we suggest a novel neural network Poisson process model which: (i) flexibly learns the connections between environmental stimuli and neural representations, and between neural representations and behavioural responses; (ii) jointly fits both behavioural and neural data; (iii) handles variabilities between response times in different trials of an experiments by a temporal rescaling mechanism, and (iv) derives spike count statistics disentangled from chosen temporal bin sizes. The framework allows efficient training of the model without making assumptions about the functional form of the relationship between input stimuli and neural and behavioural processes. We apply the method to two neural/behavioural datasets concerning visual discrimination tasks: one collected using Neuropixel probes [Steinmetz et al. 2019] from mice, and the other the output of a hierarchical network model with reciprocally connected sensory and integration circuits that was designed to model behaviour in a motion-based task [Wimmer et al. 2015]. We show that our method is able in both cases to link behavioural data with their underlying neural processes and input stimuli; the synthetic dataset allows us to compare our results against ground truth.

## 2 The model

### Data description

We model canonical visual discrimination experiments. In our case, on each trial, subjects are presented with a stimulus and have to choose an option (or keep still; NoGo). We consider two datasets in this setting. The first is the visual discrimination experiment of [Steinmetz et al. 2019] (Figure 1a). On each trial, mice are presented with a stimulus (visual contrast on the left or right side) and have to make a simple response by turning a wheel left or right or keeping it still. The second dataset is synthetic and based on the work of [Wimmer et al. 2015] (Figure 1b). A hierarchical spiking neural network model is built to capture the essence of evidence integration and decision-making of monkeys in a standard two-alternative forced-choice motion discrimination task [Gold and Shadlen 2007; Britten et al. 1996].

**Figure 1:**
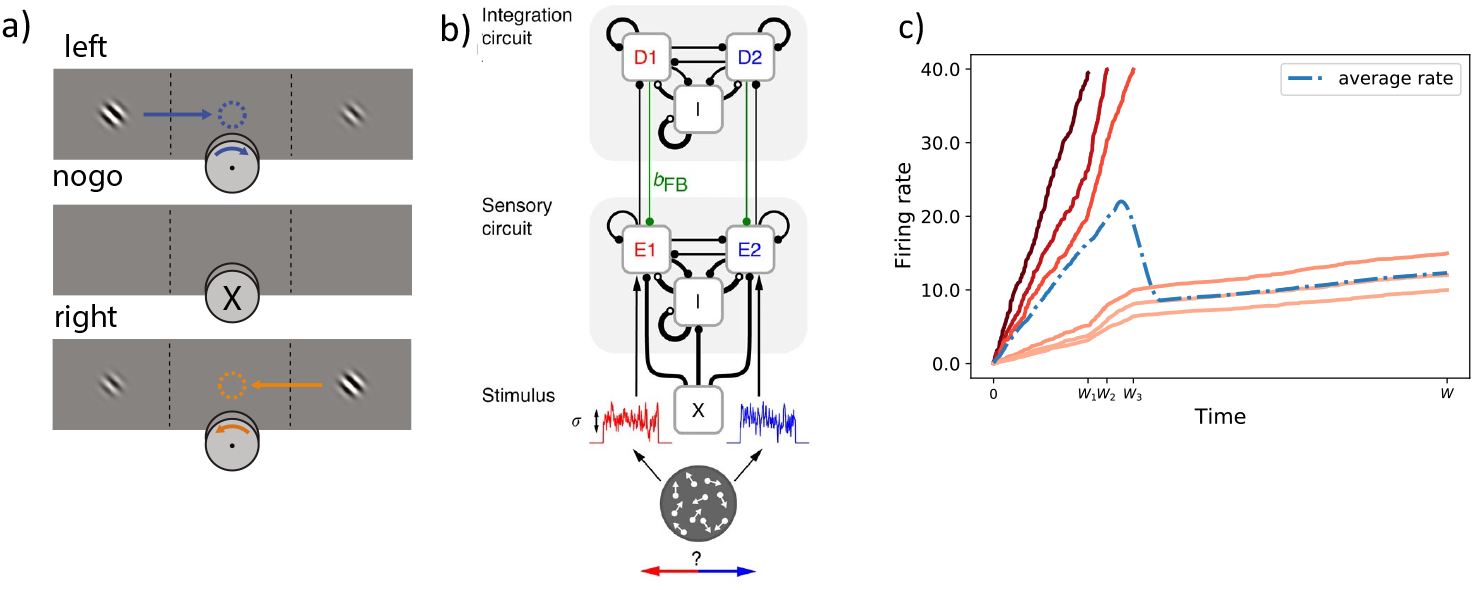
**a)** Steinmetz’ visual discrimination task [Steinmetz et al. 2019]. On each trial, visual stimuli of different contrasts were potentially presented on the left and right sides. If both sides had zero contrast, the mouse earned reward by NoGo. If both had equal, non-zero, contrast, it was rewarded at random. Otherwise, it was rewarded for reporting (*a* ∈ {left, right}) which contrast was larger. The figure is adapted from [Steinmetz et al. 2019]. **b)** Synthetic network model illustrating the sensory (E1; E2) and integrator (D1; D2) circuits enjoying feed-forward and top-down feedback connections as well as lateral excitatory and inhibitory (population I) recurrent connections within each circuit. The figure is adapted from [Wimmer et al. 2015]. **c)** Illustrative example illustrating the occurrence of systematic bias in firing rate estimation when aggregating trials with different end times. Red curves show firing rates for six sample trials. The dashed line shows the average of firing rates based on the unfinished trials at each point in time. Looking at the blue curve, one might conclude that neural activities increase and then decrease over times, which is not true for any individual trial.

### Formalisation

The total number of trials in an experiment is denoted by 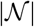; the stimuli on trial 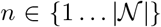 are (generically) denoted by vector **x**_*n*_. If a response was made on trial *n* (at a time relative to stimulus onset we call *r_n_*), we denote it by 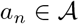 (where, here, 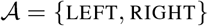. After an observation window of *W* = 400ms (for the Steinmetz dataset; same as the window size chosen in [Steinmetz et al. 2019]) and *W* = 2000ms (for the synthetic data) expires, then we consider the subject to have chosen NoGo.

Different neurons in different areas and even different animals (in the Steinmetz dataset) may be recorded and contribute in separate trials. In total, 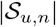 spikes are observed from unit *u* at times 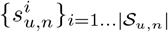, relative to stimulus onset in the corresponding trials.

In the following, we first discuss how we model the neural data; and then how we couple this model to predict behaviour. Figure 2 provides an overview of the designed framework. For more details on the model architecture, please refer to Supplementary Materials.

**Figure 2:**
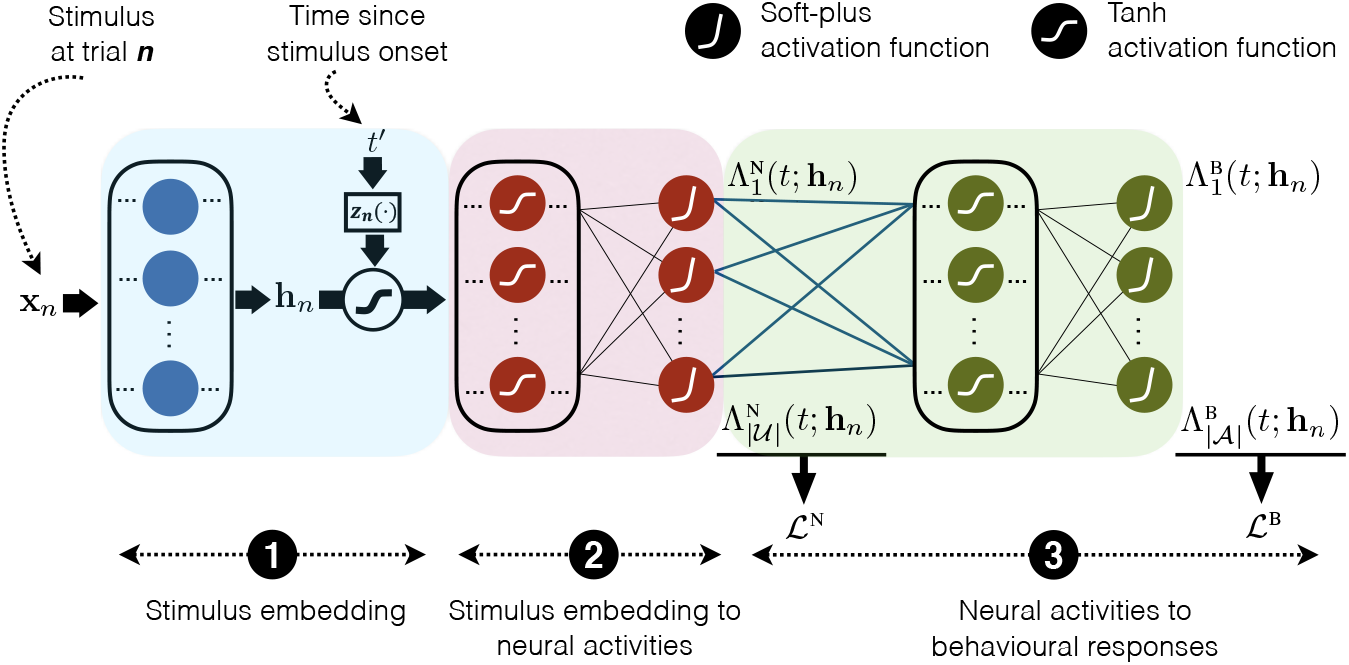
Network architecture. ① Embedding of input stimulus into vector **h**_*n*_. ② Transformation of embedding **h**_*n*_ into neural activity of each region for each time point *t* since the stimulus onset. The monotonic transformation function, *z_n_* (*t′*) = *t*., is applied to the input spike time series is step. Neural activities are characterised by the rescaled *cumulative* intensity function of spike train for each region, denoted by 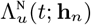 for regions 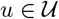. The intensity function 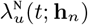 is obtained by differentiating the cumulative intensity hunction 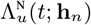 with respect to *t*. Component ② structurally ensures that 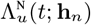 increases with time, so 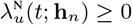. ③ Neural activities are mapped to behavioural responses which are represented by the rescaled cumulative intensity function 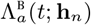 for making each action 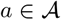 at each time *t* since stimulus onset (*t*). The data likelihood is computed over both recorded spike trains and behavioural reaponses. This yields a neural log-likelihood function 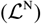 and a behavioural log-likelihood function 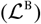.

### 2.1 Spike train models

We make the simplification that the spikes of each neuron *u* in trial *n* can be modelled as the output of an inhomogeneous Poisson process [Daley and Vere-Jones 2006] with a latent intensity function 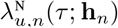 (the superscript N indicates that the intensity function is for the Neural data). This can be interpreted as the instantaneous probability of observing a spike at time *τ*, where **h**_*n*_ is a function of the stimulus **x**_*n*_. The Poisson process assumption is that successive spike times are independent, given 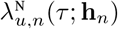. Note that we seek to capture signal correlations but not noise correlations, and so **h**n does not depend on the spikes observed during a trial.

#### 2.1.1 Time rescaling of spike trains

We consider spikes from neuron *u* until either a response *a_n_* was made at time *r_n_*, or to the end of time window *W*, whichever comes first, i.e., up to *W_n_* = min(*W, r_n_*). The reason for restricting the observation period to *r_n_* is because the aim is to model the neural processes that lead to behavioural responses, rather than what happens post-response. The joint probability density of observing spike trains from neuron *u* in trial *n* is then,

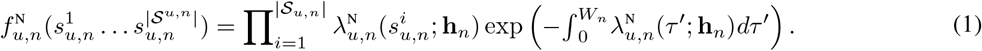

Intuitively, the term 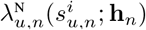 represents the probability density of observing a spike at time 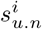. and the exponential term represents the probability of not observing spikes at other times in the observation period. We aim to estimate a single function 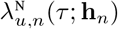 to model the neural activities across all trials. However, note that the duration of trials can be different (based on response times) and only trials that ended *after τ* can contribute to the estimation of 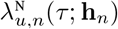, which means that this quantity is implicitly conditioned on *r_n_* > *τ*. This property makes the interpretation of 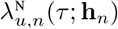 rather inconvenient since it will no longer represent how neural activities evolve over time, but is confounded by the distribution of response times (see Fig 1c). To address this issue, it is tempting to merely condition 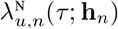 on response times (in addition to **h**_*n*_) to get a picture of the spike trains that lead to each specific response time (rather than all the response times after *τ*). This, however, only partially addresses the issue. To address it more fully, we aim to map all trials with different duration to the same time span. To achieve this goal, we propose the following theorem and proposition:

#### Theorem 1.

*Let* 0 < *s*^′1^ < *s*^′2^ <,…,< *s*^′*j*^ ≤ *W_n_* ≤ *W be a realization from an inhomogeneous Poisson point process, n, with an intensity function* λ_*n*_(*t*′) *satisfying* 0 < λ_*n*_(*t*′) *for all t*′ ∈ (0, *W_n_*]. *Define a one-to-one monotonic transformation function, where*:

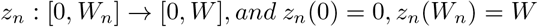

*Assume* 0 < *s*^1^ < *s*^2^ <,…, < *s^j^* ≤ *W where* ∀*k* ∈ {1,…, *j*}; *s^k^* = *z_n_* (*s*′^*k*^). *Then s^k^ are a realization from a second inhomogeneous Poisson point process with* λ(*t*) = λ_*n*_(*t*′) *where t* = *z_n_*(*t*′).

#### Proposition 1.

*For a linear transformation function z_n_*(.), *as defined in* **Theorem 1.**, *the cumulative intensity function of the original and second point process realizations are related as*: 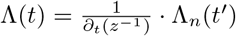.

Note that **Theorem 1.** is a special case under the Mapping Theorem [Grimmett and Stirzaker 2020]. Please see Supplementary Materials for proofs.

#### 2.1.2 Parametrising the intensity function

Here, we define 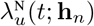 to represent canonical neural activities (prior to the response) defined over *t* ∈ [0, *W*]. Based on the above theorem, the neural activities for a certain trial with duration *W_n_* can be obtained by applying the time rescaling on the original spike time series using a monotonic function *z_n_*: [0, *W_n_*] → [0, *W*], *t* = *z_n_*(*t*′),

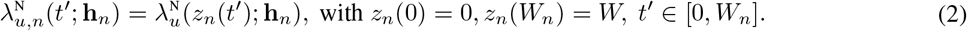

The dependence of the intensity function on the embedding **h**_*n*_ plays a crucial role in determining how neural activities are shaped by the stimulus and has to be characterized in a flexible manner. To achieve this, one option is to use a multi-layer feed-forward network which takes *t* and **h**_*n*_ as inputs, and outputs 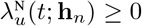. Unfortunately, it is then intractable to calculate the integral in equation 1. An elegant solution to this problem is to parameterize the cumulative intensity function 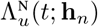 instead of 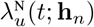 [Omi, Aihara, et al. 2019]. This is

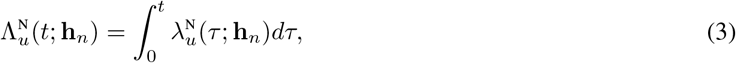

and can be (automatically) differentiated to produce the intensity function:

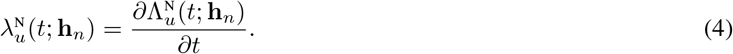

We represent 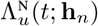 using a feed-forward network. Note that the gradient of the cumulative intensity function w.r.t to *t* must always be non-negative (see Section Model structure). In this work, we chose time to be rescaled uniformly using *z_n_*(*t*′) = *t*′*W/W_n_*. This makes a substantive assumption that the activities in slow and fast trials are stretched versions of each other. Its advantage is to reduce the task of learning trial-specific intensities to learning the canonical intensity function 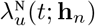. Based on Theorem 1. and Proposition 1., trial specific intensity and cumulative intensity functions (defined on *t*′ ∈ [0, *W_n_*]) are then given by,

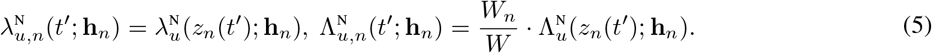

The relatively simple form of transformed cumulative intensities is a consequence of the uniform time rescaling (see Supplementary Materials for proof). Using these two related functions, the data log-likelihood implied by equation 1 is

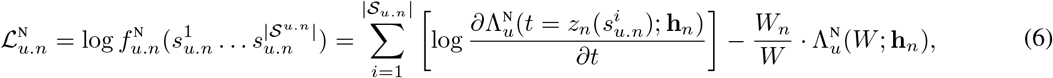

which retains the required flexibility, while obviating the calculation of the intractable integral. The total data likelihood for all the trials and neurons is then

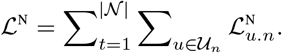

In principle, each recorded neuron in the experiment could be assigned a separate intensity function. However, given the experimental methodology of only recording some neurons on some trials in the Steinmetz dataset, the problem of missing data would be radically acute, and the model would be uninterpretable. The computational cost would also be prohibitive. Instead, we make the simplifying assumptions that all the neurons in each of the 42 brain regions identified by [Steinmetz et al. 2019] and (as is true by design) the 4 regions in the synthetic model share common intensity functions. As such, we consider 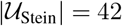 and 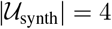.

### 2.2 Behavioural response models

Having provided a way of characterizing the neural response to the embedding **h**_*n*_, we next need to model the link between neural representations and behaviour. We assume that the probability of making a behavioural response at each point in time depends on the activity of the neurons at that time. In turn, these are driven by the stimulus **h**_*n*_ on that trial. That is, the behavioural responses are indirectly affected by the stimulus via neural activities. However, rather than model this dependence explicitly, which is hard given the punctate nature of the response, we approximate it implicitly, via smooth intensity functions that in turn depend on Λ^N^.

The intensity function for an action *a* is denoted by 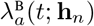, which specifies the instantaneous probability of taking the action at time *t* relative to stimulus onset. The superscript B indicates that the intensity function is for the Behavioural data. A key simplification is to allow for the theoretical possibility that the animal performs the same action more than once on a trial; or performs both actions. However, that actions are actually sparse implies that this approximation is not too costly. We write the canonical behavioural cumulative intensity function as a function of the canonical neural cumulative intensity functions.

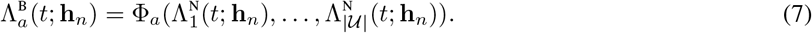

Function Φ_*a*_(.) can be realised using a deep feed-forward network which can represent arbitrary dependencies between neural activities and behavioural responses.

Then, differentiating:

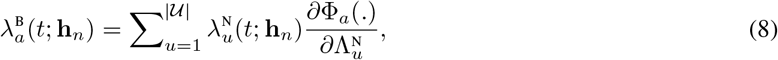

which implies that the behavioural response probability at each point in time is indirectly dependent on the stimulus through the spike rate of different neural activities, as desired. Function Φ(.) is also designed to be increasing in *t* to ensure that 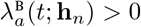 (see Section Model structure).

These response rates are presented as a function of canonical neural intensity functions. Similar to 5 for each trial *n* we have,

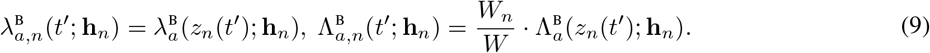

The subjects can either act or not on a trial; the latter is determined by a censoring window *W*. Write 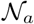 as the set of trials on which action a was taken before *W*, at reaction time *r_n_*. The simplified joint probability distribution of the behavioural observations is then:

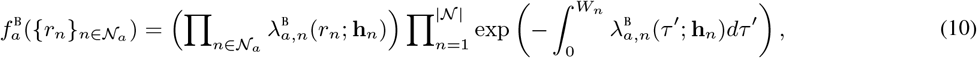

and, taking logs, the log-likelihood for those observations is,

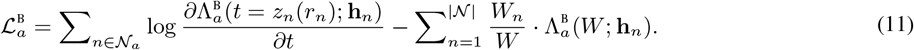

Over the whole experiment and actions, the behavioural likelihood can be defined as,

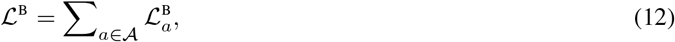

in which 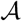 is the set of available actions.

### 2.3 Model structure

We implement the model using the neural network architecture shown in Figure 2. This has three components. The first maps the stimulus **x**_*n*_ that was presented through a series of fully connected layers to realize an input embedding denoted by **h**_*n*_. The second component takes the embedding **h**_*n*_ and *t* and outputs the modelled activity of each neural region *u* at time *t* in the form of cumulative intensity functions for 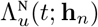. This component is designed such that the outputs of the network, i.e., 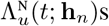, are monotonic functions of *t* to ensure that their gradients with respect to *t* (which are neural intensity functions) are always positive. To achieve this, following ideas from [Sill 1998; Chilinski and Silva 2018; Omi, Aihara, et al. 2019], the weights of the network are constrained to be positive and tanh activation functions are used in the middle layers and soft-plus in the output layers.

The third component of the model takes the neural cumulative intensity functions and maps them to the behavioural cumulative intensity functions (function Φ in equation 7). We used the same method as for the second component to ensure that the gradient of 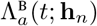 with respect to *t* is positive.

For training the model, the neural loss function 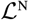 is used to train all the weights from stimulus to neural cumulative intensity functions (blue and red rectangles in Figure 2). Given these trained neural cumulative intensity functions, then the weights connecting neural outputs to behavioural outputs are trained using 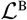. The gradients were obtained using automatic differentiation in Tensorflow [Abadi et al. 2015]. See Supplementary Materials for details.

## 3 Results

### 3.1 Data structure

#### Synthetic dataset

For the synthetic data, we use data generated from the model introduced in [Wimmer et al. 2015]. Activities from two direction-selective sensory regions (E1 and E2; e.g. V5/MT) as well as two integrator regions (D1 and D2; e.g. LIP, FEF) are modeled and then observed. Each region has 240 neurons and the whole experiment consists of 1800 trials (1200 trials for training and 600 trials for testing). Left and right sensory regions prefer leftwards and rightwards motion respectively; time-varying activity in these regions inspired by stimuli with coherence levels varying from completely obscure: 0%, to rather definite: 50% or 80% (which are encoded using one-hot encoding), are accumulated by populations in the integration region. The latter has attractor dynamics; a response is realized when the activity state is sufficiently close to one of the two attractors. A choice is modelled as being made when the average activity of a (trial-)random subset of neurons in D1 or D2 over a window of 50ms reaches 40Hz. This corresponds to strong evidence in favour of motion in the corresponding direction which can be right (for D1) or left (for D2).

#### Steinmetz dataset

We use the data reported in [Steinmetz et al. 2019]^2^. The experiments consist of 38 sessions of a visual discrimination task (Figure 1a). Activities from 42 brain regions from the left hemisphere of the brain were recorded (not all regions were recorded in all the sessions). Overall the data from 30,000 neurons were recorded and the whole experiment consisted of 10011 trials. We used the data from 12 sessions for testing and the rest for training the model.

On each trial, the animals were presented with stimuli on the left and right and were required to turn a wheel LEFT, right or keep it fixed (NoGo), based on the contrast input (four possible levels of contrast on each side: 0, 25, 50, 100%; 0% on both sides requires NoGo). We encoded stimulus contrast using one-hot encoding based on which side had a higher contrast, or whether they had equal contrasts (**x**_*n*_ of dimension 3). The reaction time *r_*n*_* corresponds to the beginning of the wheel turn if this happens before the end of the response window.

### 3.2 Training process

All the weights in the model were trained using the Adam optimiser [Kingma and Ba 2014]. Stimulus and integrator components (blue and red rectangles in Figure 2) each were composed of three fully connected hidden layers with 20 neurons in each layer. The third layer was then connected to the output layer consisting of one neuron per region in the dataset. The neural component was followed by two fully connected behavioural layers for each action (with 10 neurons). ‘softplus’ and ‘tanh’ activation functions were used to ensure positive of intensity functions. See Supplementary Materials for more details about the model architecture and training process.

### 3.3 Experiments

We show statistics of the quality of the fit of the model later. However, we first illustrate the neural and behavioural properties of the model by freezing the weights and performing simulations with different values of Wn for sample regions of both synthetic and Steinmetz datasets. Results showing the performance of the model on all the available test regions of both datasets are presented in Supplementary Materials.

The solid lines in the upper panels of Figure 3 illustrate the neural responses for the synthetic dataset for 0.3 ≤ *W_n_* ≤ 0.6 when the stimulus had highest coherence level (0.8) and moved right. The chosen interval includes more than 90% of all trials in the coherence level of 0.8. These results are compared with the empirical activity derived from the data (dashed lines) for both integrator regions D1 and D2. Note the change in *y*-scale between the plots, given the strong left stimulus. These values are closely related to the learned intensities and capture the observed variability in response times between different trials given the time rescaling of input spike trains.

**Figure 3:**
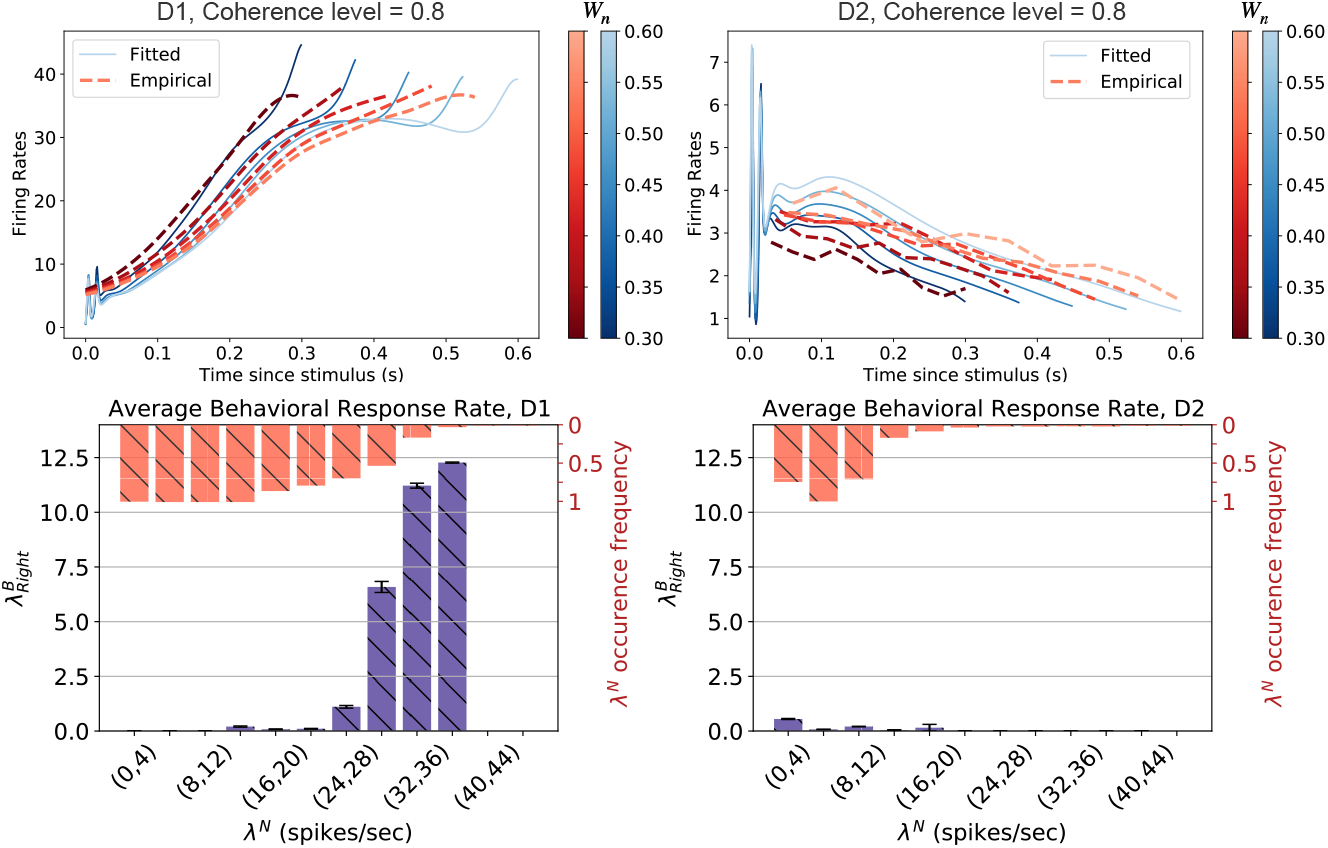
Synthetic Data (Upper panels). The activity rate of neural population in the two integrator regions of the synthetic dataset. The dashed lines show the empirically derived firing rates compared to the solid lines which are estimated using the proposed model. Each line plot corresponds to the average activity rate of trials with *W_n_* in a specific interval illustrated by the colorbar (see section F in Supplementary Materials). **(Lower panels).** Illustrating the average response rate for each interval of neural activity rates. The plots correspond to the trials with highest coherence level. The plots show the average between trials with a right reaction. Error bars show the standard error. Purple bars show the average behavioural response rates. Orange bars indicate the proportion of trials with the occurrence of each neural response interval.

Figure 3 lower panels show the mean rate of behavioural responses for each interval of learned neural intensities for the complete set of trials. As expected, the highest rate of behavioural action is observed for neural intensities close to 40 spikes/sec for D1. By contrast, region D2 shows very low behavioural activity rates due to the non-favorable direction of stimulus in these trials. Note that the shown empirical firing rates are averaged across all responses, while the intensities correspond to different response times, which helps explain the differences between the two measures. For example, in very fast responses, there is an initial burst in the firing rates, which is captured by the initial sharp rise in the intensities (see Figure 3), but this is invisible in the averaged firing rates.

The upper panels of Figure 4 show modelled and actual neural activities in the Steinmetz dataset for two example brain regions with high contrast levels of stimulus on right: one the subiculum (sub), which [Steinmetz et al. 2019] reported as containing neurons that consistently fired before wheel turns regardless of their direction (arguably a surprising feature of this dataset, given the relationship of the subiculum with areas such as the hippocampus and entorhinal cortex rather than motor regions), and the other, visual (visp; primary visual area), which is reported to have the highest portion of visual encoding neurons. The activities were all recorded in the left hemisphere, and therefore, the right stimulus/action are contralateral to the recording sites. Comparing the lower and upper panels of Figure 4 show, we generally see more activity on the left side in visp for stimuli with high contrast levels on the right which is consistent with previous analyses [Steinmetz et al. 2019]. It is clear that the region shows lower neural activity levels for ipsilateral stimuli. The subiculum (sub), is reported in [Steinmetz et al. 2019] as containing neurons that consistently fired before wheel turns regardless of their direction. We can see in Figure 4 lower panels that this indeed is captured by our model for the SUB region where it reaches similar firing rates no matter the direction of stimulus and motion are. The relative timing of the activities in the areas are also consistent with expectations. The solid lines in the panels show estimated firing rates, which are consistent with the data (dashed lines) in particular for visp which had a peak in firing rate early after stimulus was presented (see below) and closely related to the presented visual stimuli in the task. Note that in our framework, estimation of the neural activities (firing rates) does not rely on selecting a temporal bin size. This is unlike most previous state-of-the-art works [Liu and Lengyel 2021] where the output firing rates are substantially affected by the choice of bin size.

**Figure 4:**
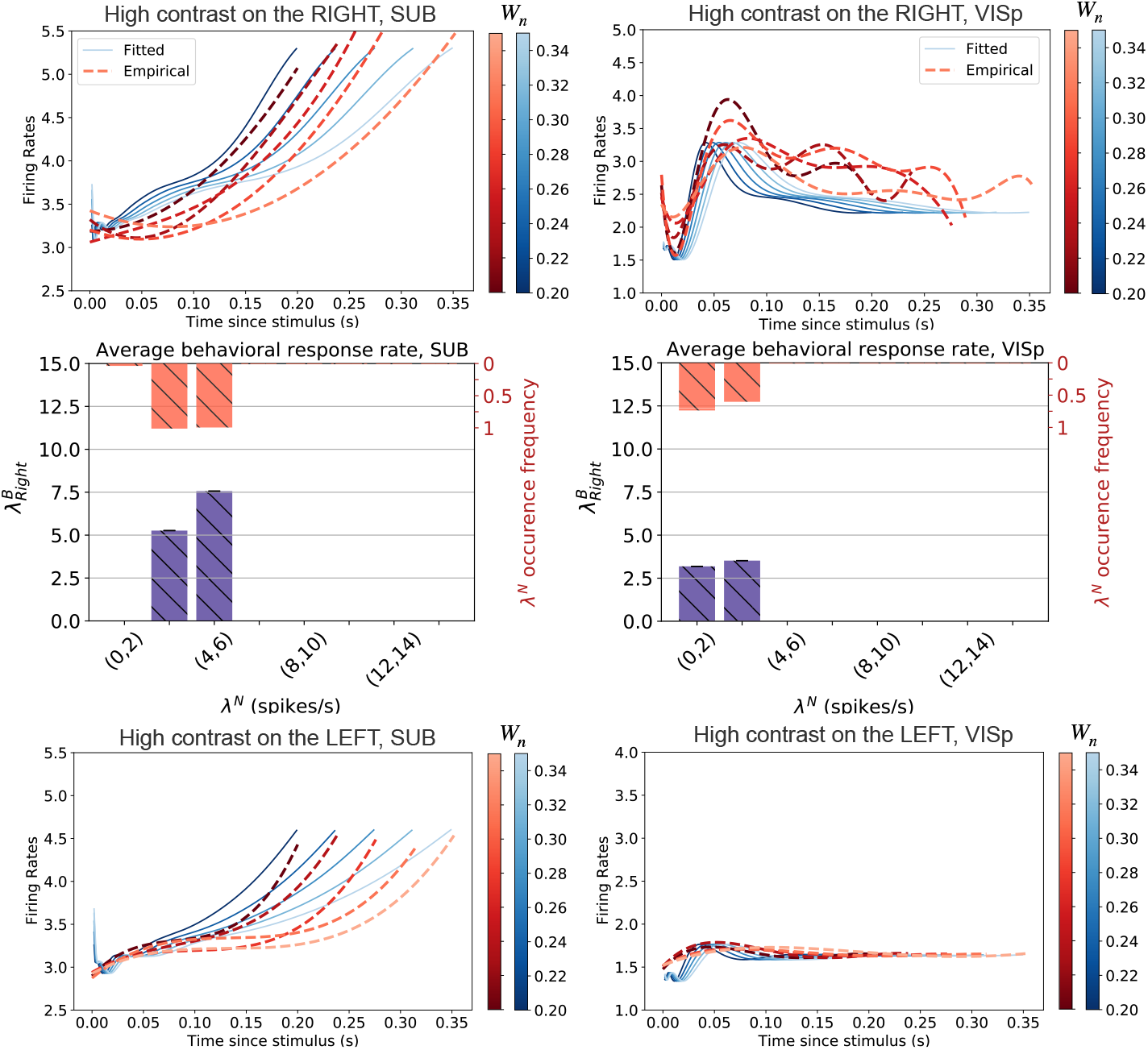
Steinmetz (Upper panels). The activity rate of neural population in the two sample regions of the Steinmetz dataset for the highest contrast level on right. The dashed lines show the empirically derived firing rates compared to the solid lines which are estimated using the proposed model. Each line plot corresponds to the average activity rate of trials with *W_n_* in a specific interval illustrated by the colorbar (see section F in Supplementary Materials). **(Middle panels).** Illustrating the average response rate for each interval of neural activity rates. The plots correspond to the trials with high contrast on right. Error bars show the standard error. Purple bars show the average behavioural response rates. Orange bars indicate the proportion of trials with the occurrence of each neural response interval. **(Lower panels).** Neural activities for contralateral and ipsilateral stimuli. The visp shows a direction selective activity pattern, preferring the contralateral stimuli on the right. However this is not the case for SUB which is in agreement with reports in [Steinmetz et al. 2019] where this region is known to contain firing neurons before the motion initiation regardless of the direction of stimulus and movement.

Next, examining the behavioural predictions of our model, the neural activities presented in Figure 4 show that sooner responses (with high probabilities) are strongly related to the peak intensities in visp. In agreement with these findings, middle right panel in Figure 4 shows higher probability for the occurrence of reactions when the neural activity in visp peaks. sub is also coupled to behaviour in the middle panel – in fact for both directions of movement.

### 3.4 Baseline Methods

Finally, Table 1 shows a comparison of the performance of our proposed framework to those of recent prominent baseline point process estimators. We compare the negative log likelihoods (NLL) for the neural activity on sample regions of Steinmetz dataset as well as the 4 regions of the synthetic dataset. For details on the utilized settings for implementing the baseline methods, please see the Supplementary Materials.

- **GLM Model [Truccolo et al. 2005; Paninski 2004]**: Generalized-linear models (GLM) also known as Poisson regression, are used to model the intensity of the input data as a linear combination of time-dependent covariates. Here, the total spike counts for all trials are calculated and concatenated for count windows of 5ms as inputs to the GLM model.
- **NHPoisson Model [Cebrian 2015]**: NHPoisson is a method for the modelling non homogeneous Poisson processes in time estimating maximum likelihood. The model is based on formulating the intensity as a function of time-dependent covariates.
- **Universal Count Model [Liu and Lengyel 2021]**: This model builds on sparse Gaussian processes (GP) to capture arbitrary spike count distributions flexibly relying on both observed and latent covariates. It uses scalable variational inference and can jointly infer the covariate-to-spike count distribution mappings and latent trajectories. We also examine a second variant of this model which replaces the GP-based approaches with an artificial neural network (ANN) mapping. We denote the two variations by U-GP and U-ANN respectively.
- **Poisson Gaussian-Process Latent Variable Model (P-GPL) [Wu et al. 2017]**: In this model, Poisson spiking observations are accompanied by two underlying Gaussian processes: One governing a temporal latent variable, the other governing a set of nonlinear tuning curves. The model learns using a decoupled Laplace approximation which is a fast approximate inference method. The same set of temporal covariates as above are also utilized in the implementation of this method.

**Table 1:** Comparison of the Negative Log-Likelihood (NLL) measure for the neural intensity function estimations in example regions of the synthetic and Steinmetz datasets.

| Methods | Synthetic Dataset |  |  |  | Steinmetz Dataset |  |  |  |
| --- | --- | --- | --- | --- | --- | --- | --- | --- |
|  | D1 | D2 | E1 | E2 | VISp | SUB | VISam | SNr |
| NN-Poisson | **-5285.118** | **-2703.695** | **-14708.950** | **-8570.335** | **3.451** | **-2.522** | **0.022** | **-19.217** |
| GLM | **-324.343** | **-146.556** | **-439.662** | **-347.569** | **87.114** | **17.219** | **2.216** | **-5.782** |
| NHPoisson | **-1330.065** | **-1066.044** | **-1354.954** | **-1341.548** | **33.817** | **-0.012** | **2.915** | **-11.871** |
| U-GP | **-3532.831** | **-2317.245** | **-10334.716** | **-5289.174** | **12.337** | **1.336** | **1.066** | **-17.484** |
| U-ANN | **-3417.905** | **-2010.752** | **-4981.384** | **-2863.996** | **12.679** | **1.354** | **1.108** | **-14.603** |
| P-GPL | **-2631.857** | **-2196.593** | **-4774.682** | **-3729.311** | **12.375** | **-1.312** | **1.584** | **-15.173** |

It is important to mention that the performance of all the baseline methods depend on the length of the selected time bin for spike count calculation; a constraining dependency causing non-robust results that our proposed method overcomes. We selected the time bin length for optimal performance by performing a heuristic search in a rational range.

Note that the average spike counts per region in the Steinmetz dataset is roughly 60 times less than in the synthetic one. This is due to the difference between the lengths of the experiment and the numbers of neurons per region in the Steinmetz and synthetic dataset. Thereby, the NLL measures differ by two orders of magnitude.

Finally, Figure 5 shows the comparison of the performance of the proposed model with the baseline methods in estimating neural activities summed over all the 37 regions in the test data of Steinmetz dataset (including the 4 regions reported in Table 1).

**Figure 5:**
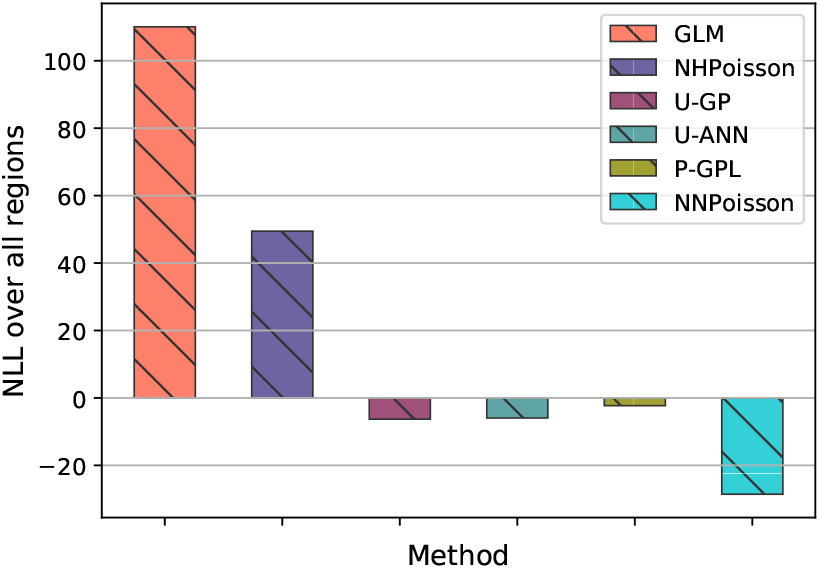
Total NLL of the estimated neural intensity function on Steinmetz test set. Illustrating the sum of NLL values evaluated over all the 37 test regions in the Steinmetz dataset using different baselines compared to the performance of the proposed method.

## 4 Discussion

We presented a novel framework for linking neural spike trains to sensory inputs and behaviour. The framework extended previous works on fMRI data [Dezfouli et al. 2018] by using a flexible Point process framework. The model was able to learn a suitable encoding of the stimulus and provided a joint explanation for both behavioural and neural data that could be used to recover correlational links between neural and behavioural activities. Unlike previous efforts, the learning process of the proposed model is independent of the selection of a time bin for spike count calculations obtaining higher robustness. The current method represents the dependency of neural activities on stimulus and trial duration, but not on previous neural activities – thus capturing signal rather than noise correlations (although the latter are an obvious target for future work). There are many additional directions for future work: capturing richer aspects of behaviour that are known to couple to neural activity [Balleine and O’Doherty 2010]; integrating and/or substituting spiking activity with calcium imaging; like auto-regressive linear-nonlinear-Poisson (LNP) models [Chichilnisky 2001]; differentiating more finely the activity in different regions (including neurons with opposite stimulus coding). Nevertheless, we suggest that our method casts brain and behaviour interactions in a compelling new light.

## Supplementary Materials

### A Proof of Theorem 1

Following the Mapping Theorem of an inhomogeneous Poisson process [Grimmett and Stirzaker 2020], and given a one-to-one monotonic transformation function, *z_n_*(*t*′) = *t*, between the original inhomogeneous Poisson point process 0 < *s*′^1^ < *s*′^2^ <,…,< *s*′^j^ ≤ *W_n_* ≤ *W* and 0 < *s*^1^ < *s*^2^ <,…, < *s^j^* ≤ *W*, let 0 < *s*^1^ < *s*^2^ <,…, < *s^j^* ≤ *W* represent a set of event (spike) times from a second inhomogeneous Poisson point process. Let *g*(*t*) represent the corresponding event time probability density. Given *g*(*t*) is a density function (measurable and non-negative function), following the push-forward probability density, we get:

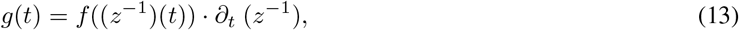

where *z*^-1^ is the inverse of the transformation, *z_n_*(.), and *f* (*t*′) is the event time probability density function of the original Poisson point process.

Now, for *t* ∈ (0, *W*], let *N*(*t*) be the sample path of the associated counting process. The sample path is a right continuous function that jumps 1 at the event times and is constant otherwise. Then, we compute the probability that a spike *s^k^* occurs in [*t, t*+Δ*t*) where *k* = *N*(*t*) + 1. Note that events {*N*(*t*+Δ*t*)–*N*(*t*) = 1} and {*s^k^* < *t*+Δ*t* |*s^k^* > *t*} are equivalent. Thereby,

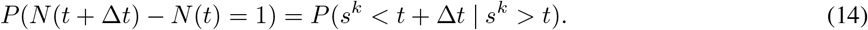

From the definition of conditional probability:

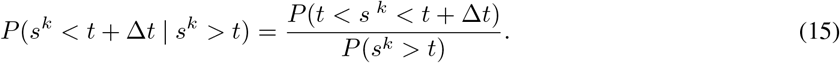

Therefore, we get:

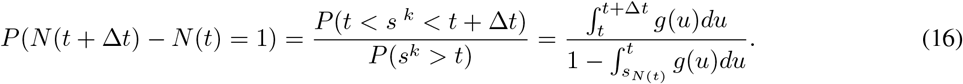

Using equation 13, we obtain:

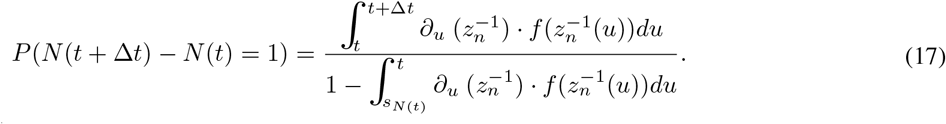

We also have 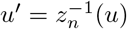 and hence:

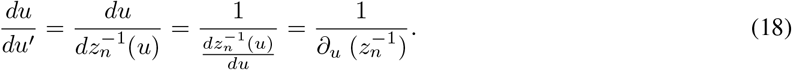

Thereby, inserting the above in equation 17 and by a change of variable *u* to *u*′, noting that 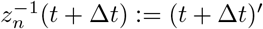, we get:

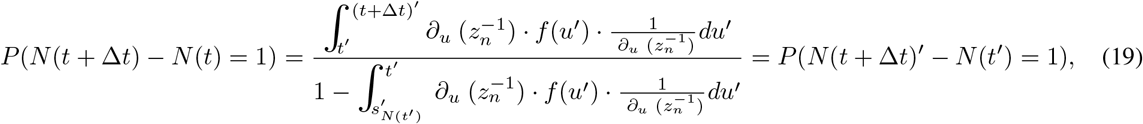

where *N*(*t*′) is the sample path of the associated counting process for *t*′ ∈ (0, *W_n_*]. The intensity function of a point process, λ(*t*), can be written as:

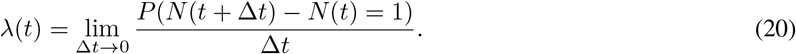

hence, since we proved *P*(*N*(*t* + Δ*t*) < *N*(*t*) = 1) = *P*(*N*(*t* + Δ*t*)′ – *N*(*t*′) = 1), we get:

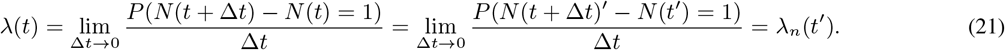

Finally, the intensity function of the new count process λ(*t*) equals λ_*n*_(*t*′) of the original Poisson point process, thereby satisfying the four properties of a Poisson point process [Ross et al. 1996] and we have now established our result. Figure 6 is schematically illustrating this result in case of a linear transformation function.

**Figure 6:**
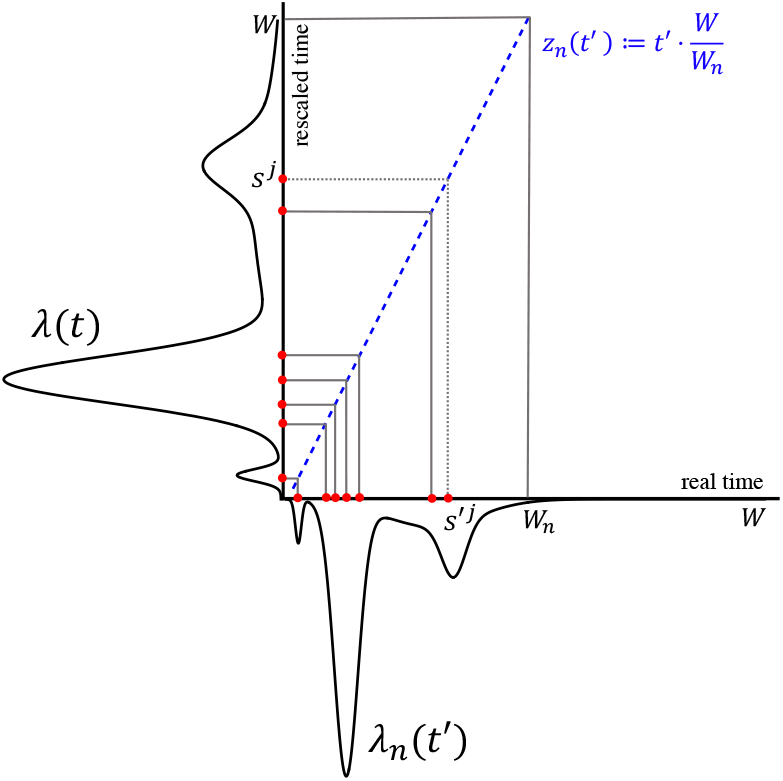
Illustrating the transformation of the Poisson point process in real time 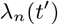, to the rescaled time domain which results a point process with intensity function λ(*t*). The transformation function 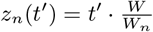 performs a one-to-one monotonic mapping of the events in real time (*s^′j^*) to the events on the rescaled time axis (*s^j^*).

**Figure 7:**
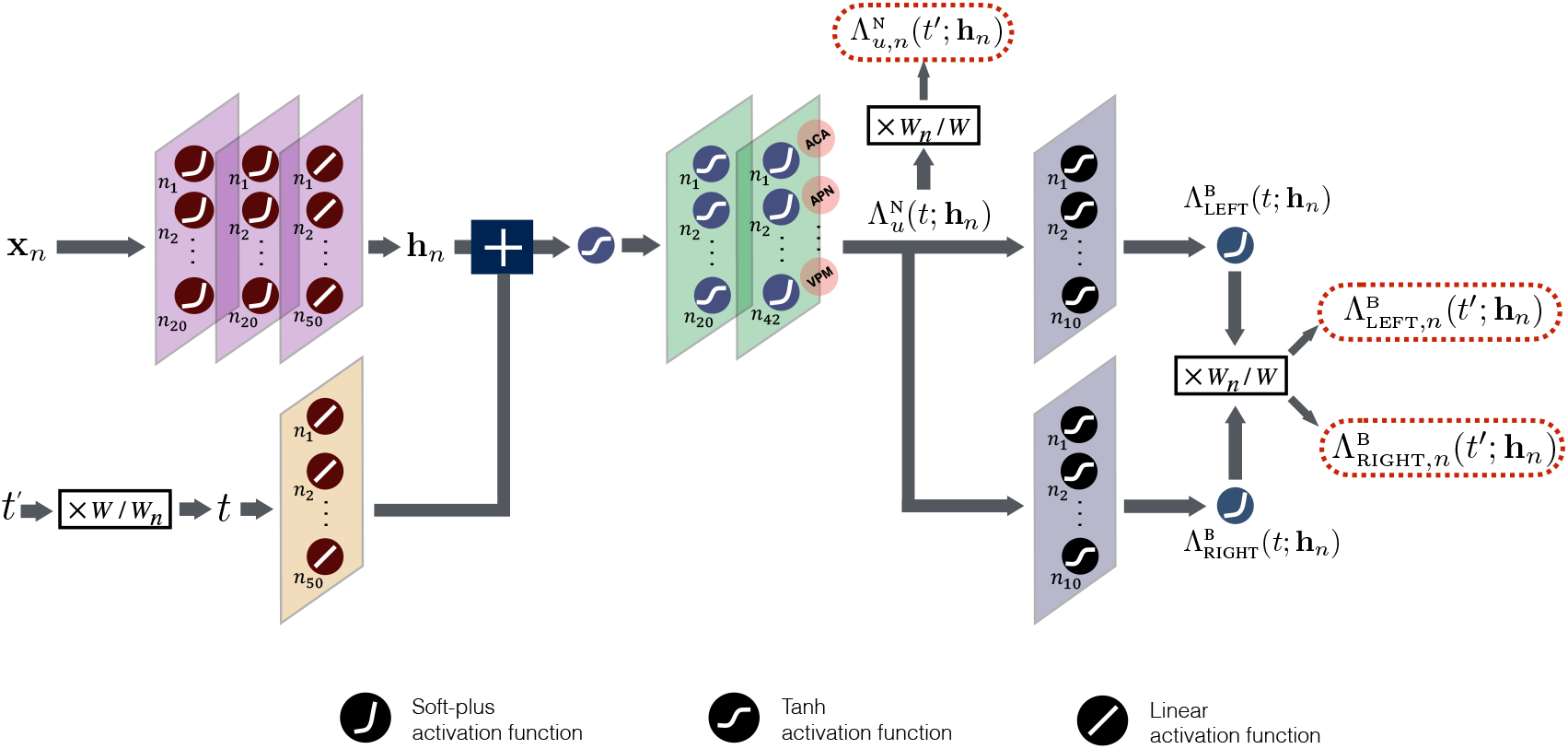
Model Architecture. Schematic representing the details of the implemented neural network to be trained on the Steinmetz dataset. The neural and behavioural outputs of this network are multiplied by a factor of *W_n_/W* for the selected linear transformation function, *z*, to obtain the final model readouts.

### B Proof of Proposition 1

The cumulative intensity function of a point process with intensity function λ_*n*_ (*t*′) is given by:

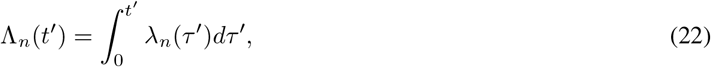

for *t*′ ∈ [0, *W_n_*], which can be (automatically) differentiated to produce the intensity function,

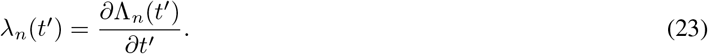

Given a monotonic transformation function *z_n_*(*t*′) = *t*, and since λ_*n*_(*t*′) = λ(*t*) (see A), we get:

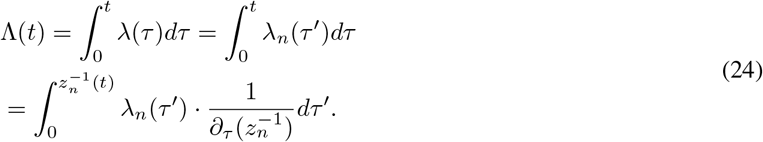

Now, following integration by parts,

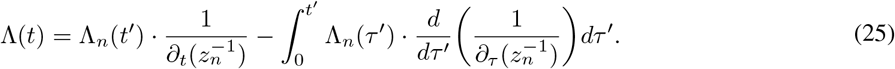

Inserting the special case of a linear transformation function where 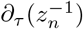 is a constant, the above equation reduces to:

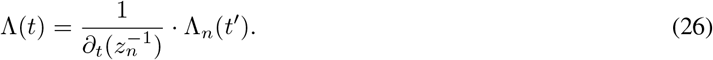

In this work, we rescaled time using function *z_n_*(*t*′) = *t*′*W/W_n_* where *z_n_*: [0, *W_n_*] → [0, *W*] and we chose to stretch the firing rates. Thereby, trial specific cumulative intensity functions are simply defined as:

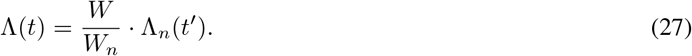

### C Model architecture and training

In this section we explain the details of the model architecture.

#### C.1 Steinmetz dataset

The input stimulus **x**_*n*_ is passed through two dense layers with 20 units (softplus activation). The output is then passed through a layer with |**h**_*n*_| =50 units using a linear activation function to create the stimulus embedding, **h**_*n*_. There is no constraint on the weights in these layers.

As mentioned in the main text, we assumed the time rescaling function is linear. The elapsed time *t*′ ∈ [0, *W_n_*] (since the stimulus) is first scaled by a factor of *W/W_n_* to obtain *t* ∈ [0, *W*] and is then passed through a linear layer (with the same number of units as |**h**_*n*_| = 50) with the weights constrained to be non-negative. The resulting output is added to **h**_*n*_, with the sum then being passed through a tanh activation function.

The output of this tanh activation is passed through a dense layer with 20 units and a further tanh activation function (first neural layer) and then through another dense layer with 42 units and softplus activation function (second neural layer). The weights of these layers are all constrained to be non-negative. The outputs of this layer correspond to 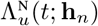, referred to as neural outputs, and are multiplied by *W_n_/W* in the network readout to obtain 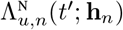 for *u* =1… 42 which is then passed to the loss function.

For modelling behavioural data, the neural output 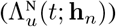 is passed through a dense layer with 10 units (tanh activation) and then through another dense layer with 1 unit (softplus activation) to produce 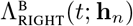. The weights are all constrained to be non-negative.

The neural output 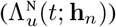 is also passed through a dense layer with 10 units (tanh activation) and then through another dense layer with 1 unit (softplus activation) to produce 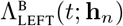. The weights of all the layers are constrained to be non-negative.

Note that the path from *t* to neural and behavioural outputs only contains positive weights which, together with monotonic activation functions, is a sufficient condition to guarantee that the outputs (neural and behavioural) are monotonic functions of *t* [Sill 1998; Chilinski and Silva 2018; Omi, Aihara, et al. 2019].

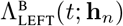 and 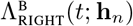 were multiplied by factor of *W_n_/W* to obtain trial specific values for 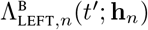 and 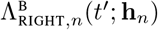.

Figure 7 visualizes the details of the model structure implemented to be trained on the Steinmetz dataset.

#### C.2 Synthetic dataset

The same model architecture was used for this dataset as explained above, but with the following differences: (i) the size of output neural layer was set to four for the four regions involved in this dataset; (ii) the size of the embedding was set to |**h**_*n*_| = 20, which was smaller than for the Steinmetz dataset, since the synthetic dataset had a smaller number of regions.

#### C.3 Training

Parameter adaptation was performed using the Adam optimizer with learning rates of 0.01 (Steinmetz) and 0.001 (synthetic). The neural loss function was used to train all the weightsup tothe neural outputs, and behavioural loss was used to train all the weights connecting neural outputs to the behavioural outputs. For the Steinmetz dataset, in each iteration of training the weights were updated using neural loss for 10 steps, and then the behavioural loss was used to update the weights for 420 steps (eac.i step is one update to the weights). The training process continued until no significant improvement op the losses (ontraining datn) was observed. For the synthetic dataset, in each iteration the neural loss was used to update the weights for 10 sieps, which was followed by training using the behavioural loss for 40 steps.

For the generation of the synthetic data we used the code provided in this link^3^. We generated 1200 trials for training and 600 trials for lesfing. For the Steinmelz datateti wrused 12 sessionsfor testing and the rest for training. Note that since in each session only a subset of regions wnre recorded (42 regions in total), we had 42 regions for training the model and 37 regions for testing the model. The split of sessions between training and test was chosen so as to obtain the maximum number of regions in bosh training and test datasets. The data was downloaded from this link^4^.

Automatic differentiation in Tensorflow was used to calculate the intensities.

### D Effects of spike time rescaling on model performance

To illustrate the effectiveness of our proposed model and the increase in model performance compared to the case where the time rescaling block is disabled, we reproduced both networks and evaluated them on the synthetic dataset. The two networks were identical in all other aspects. The NLL measures computed on the test data for both networks in all 4 regions as well as the scores of the no scaling version of the model relative to the original model are listed in Table 2; the positive score means that the proposed model (with time rescaling) is better than the case without rescaling.

**Table 2:** Comparison of the Negative Log-Likelihood (NLL) measure for the neural intensity function estimations in the original model with and without time rescaling.

| Network Model | Synthetic Dataset |  |  |  | Total |
| --- | --- | --- | --- | --- | --- |
|  | D1 | D2 | E1 | E2 |  |
| NN-Poisson (time rescaling) | **-5285.118** | **-2703.695** | **-14708.95** | **-8570.335** | **-31268.098** |
| NN-Poisson (no time rescaling) | **-5038.888** | **-2477.002** | **-14574.856** | **-8394.009** | **-30484.717** |
| Relative score | **0.04** | **0.08** | **0.009** | **0.02** | **0.025** |

### E Baseline Methods

For the comparison of our method with the available baselines, we use Generalized-linear models (GLM) [Truccolo et al. 2005; Paninski 2004] as well as a recent method for modelling non homogeneous Poisson processes (NHPoisson) [Cebrian 2015].

#### E.1 GLM

For comparison with a Poisson family GLM, one Poisson GLM with a continuous predictor (elapsed time), a categorical predictor with three/four levels for synthetic/Steinmetz dataset (stimuli coherence/contrast), as well as a behavioral predictor (the reaction times) is used to estimate the neural intensity function of each region. As the continuous predictor, we include a list of sinusoidal, tanh, and exponential functions of the elapsed time as covariates. The observed behavioral covariates are in the form of temporal arrays consisting of no-action:0, right reaction time, or -left reaction time in each time bin. Note that the sign of the provided reaction time is indicative of the direction of the motion. The total spike counts for all trials were then calculated and concatenated for count windows of 5ms. Due to the large data size and the use of aggregated spike counts from all trials, we did not use self-coupling terms for spike events. The spike counts were then used as targets for the GLM model, leading to a specific neural intensity function. The derived intensity function was then used to calculate the negative log-likelihood (NLL) based on equation 1.

#### E.2 NHPoisson

We used the same procedure as above with the count windows equal to 0.5 ms for the Steinmetz and synthetic datasets to extract the regional activity rates exploiting the elapsed time, trial stimulus type, and the behavioral reaction times. The NHPoisson R package [Cebrian 2015] requires binary information about whether a given event has occurred in each time bin. Thereby, we used a smaller count window with an order of magnitude similar to inter spike intervals. This enabled us to keep the average spike events per non empty time bin equal to 0.97 and 0.99 for Steinmetz and synthetic datasets respectively. A binary vector then stored the presence or absence of spikes in each time bin and then non-zero indices were fed to the NHPoisson model. Repeatedly, in addition to elapsed time arrays and the categorical array of stimuli coherence/contrast levels for synthetic/Steinmetz dataset, temporal arrays consisting of no-action:0, right reaction time, or -left reaction time in each time bin were also provided as behavioural covariates. We also examined a list of sinusoidal, tanh, and exponential functions of the elapsed time as covariates. Using the embedded Akaike information criterion (AIC) calculator in the package, the best covariate was selected and added to the model (lowest AIC score). The extracted neural intensity functions for each region were then obtained. The derived intensity functions were plugged into equation 1 to obtain NLL values for the NHPoisson model.

#### E.3 Universal Count Model

This universal probabilistic spike count model uses sparse Gaussian processes to derive spike count distributions. It consists of *C* Gaussian process (GP) priors, a basis expansion, and a linear-softmax mapping [Liu and Lengyel 2021]. Using 1 ms time bins, the spike counts of all trials associated with each neuron (in the Steinmetz dataset) or the neuronal population (in the synthetic dataset) are calculated and concatenated together. Note that for the case of the synthetic dataset, since the generated spike time sequences are not associated with specific neuron IDs and represent the population activity, a single cumulative spike count sequence was fed to the model for training. Once again, we use the elapsed time, the categorical stimulus type associated with each trial repeated for the duration of the trial, as well as the behavioural temporal array (consisting of 0 (i.e. NoGo), right reaction time, or left reaction time in each time bin) as the observed covariates for the model. To implement the original model (U-GP), we set hyperparameter *C* = 3 as suggested in [Liu and Lengyel 2021] and choose an elementwise linear-exponential basis expansion. We use 5 fold-cross validation on the data from each region and we cross-validate over the neuron dimension by using the train set to infer the latent states in the test data, and then evaluate the cross-validated log-likelihood of the fitted model. For the case of synthetic dataset, the cross validation is across the trials. The learning rate is set to 0.001 and we choose a tuple batch size (to indicate the trial structure of the data; for details please refer to [Liu and Lengyel 2021]) equal to the number of time bins accumulated over all trials associated with each region. In this method, an inference model with log likelihood based objectives using variational inference is built upon the data and spike couplings are computed and added as well.

In the second variation of the model (U-ANN), the same setting is implemented by replacing the GP mapping with an artificial neural network (ANN) mapping. In this model, a 10-fold cross validation is utilized. The remainder of the parameters and model settings are tuned according to the suggestions in [Liu and Lengyel 2021].

#### E.4 Poisson Gaussian-Process Latent Variable (P-GPL)

This model uses Poisson spiking observations and two underlying Gaussian processes [Wu et al. 2017]. A fast approximate inference method called the decoupled Laplace approximation is applied to learn the model from data. A Gaussian process is first used to extract the nonlinear evolution of the latent dynamic in the form of a latent variable. A second GP then generates the log of the tuning curve as a nonlinear function of the latent variable. This curve is then mapped to a final tuning curve via an exponential link function to estimate the spike rates of each neuron (λ_*i*_(*t*) for the *ith* neuron) and henceforth obtain the population activity rate in each discrete time bin. Here, the spike count matrix consisting of spike counts in time bins of 1 ms for each neuron (or the whole population in the synthetic dataset) is used to construct a generative model of the latent structure underlying these data. The data from all trials for each neuron are once again concatenated. Both Sinusoid and deterministic Gaussian bump tuning curves are examined to estimate the latent processes. The deterministic Gaussian bump tuning curve was then selected to report the results due to higher performance on our data which was expected given the naturally 2D motion space which is present in the Steinmetz dataset. We first estimated the parameters for the mapping function using spike trains from all the neurons within the training dataset. Then these parameters were fixed and the latent process using spike trains from 70% of the test data were inferred (as suggested in [Wu et al. 2017]). We then report the NLL measure comparing the estimated latent process generating λ_*i*_(*t*) values and the known empirical rates from the remaining test data averaged over all neurons in each region. The rest of the parameters are chosen according to [Wu et al. 2017].

### F Calculation of empirical rates

For the empirical firing rates, we divided the observation period into equally-sized bins each having 0.008s and 0.05s width for the Steintmetz and synthetic datasets respectively. Then, we calculated the total number of spikes in each period and normalized that by the total number of neurons and trials contributed to the spike set and also by the period duration (0.008s or 0.05s). Note that only spikes which were made *before* the response were included in the analysis. Moreover, given the selected stimulus type for each examined *W_n_* in Figure 4, only trials which had end times in an interval of 37.5 ms before the *W_n_* were included. This interval was chosen to be 60 ms for the results in Figure 3. This interval’s width was adjusted so that for each region in both datasets, at least 15% of all trials with the chosen stimulus type would fall within the interval before each of the 5 selected *W_n_*s in Figures 8 and 10.

**Figure 8:**
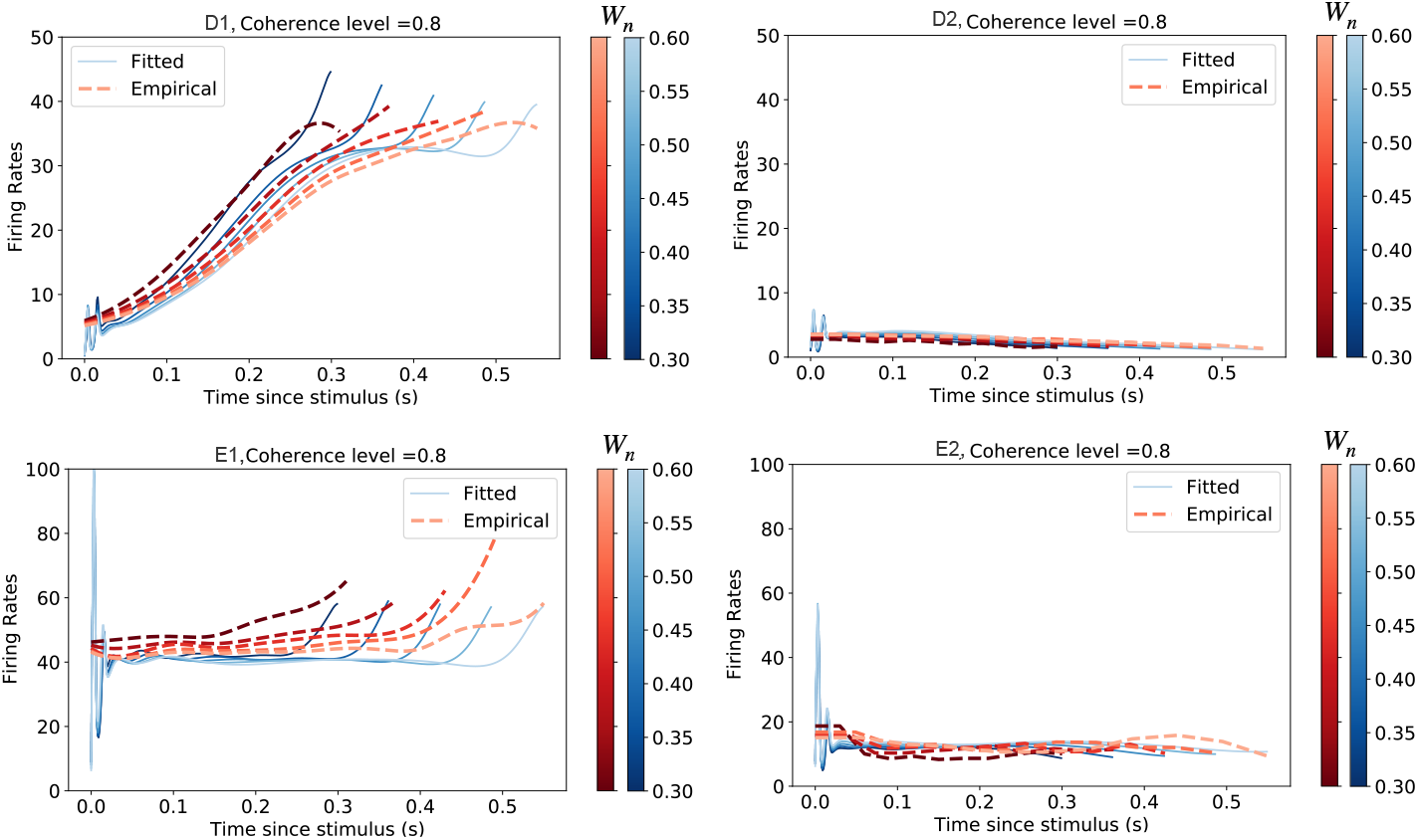
Synthetic Dataset. The red dashed lines are the empirically derived firing rates compared to the blue solid lines which are estimated using the proposed model. The plots correspond to the trials with high contrast on right. As expected from the dataset architecture, for trials when the right choice is made, the activity in D1 represents an increasing trend until the response is made. This is in contrast with region D2 for these trials where there is little to no activity detected in this region. Neurons in both sensory regions, E1 and E2 show constant activity throughout each trial with higher activity detected by E1 neurons which prefer rightward motion.

### G Additional results

#### G.1 Synthetic dataset

Using the procedure explained in Section F, Figure 8 shows the comparison of the model estimation to the empirically derived firing rates for all the regions in the synthetic dataset.

To evaluate the performance of the model in terms of predicting the behaviour of the neural population and the reaction times, we used all trials with rightward motion and fed them to the trained network of the model to get the estimated neural and behavioural response rates. The average of the estimated behavioural response rates corresponding to each of a set of potential intervals of the estimated neural responses were then calculated over trials. Figure 9 shows the results for all 4 regions. The purple bars show the average behavioural response rate in each occurred neural activity interval. The orange bars indicate the proportion of trials which achieve each specific neural activity range among all right trials, hence its occurrence frequency. Comparing the sensory regions in this figure, the behavioural correlate is with a low firing rate for E2 and with significantly higher firing rates in E1 which is the rightward selective sensory region.

**Figure 9:**
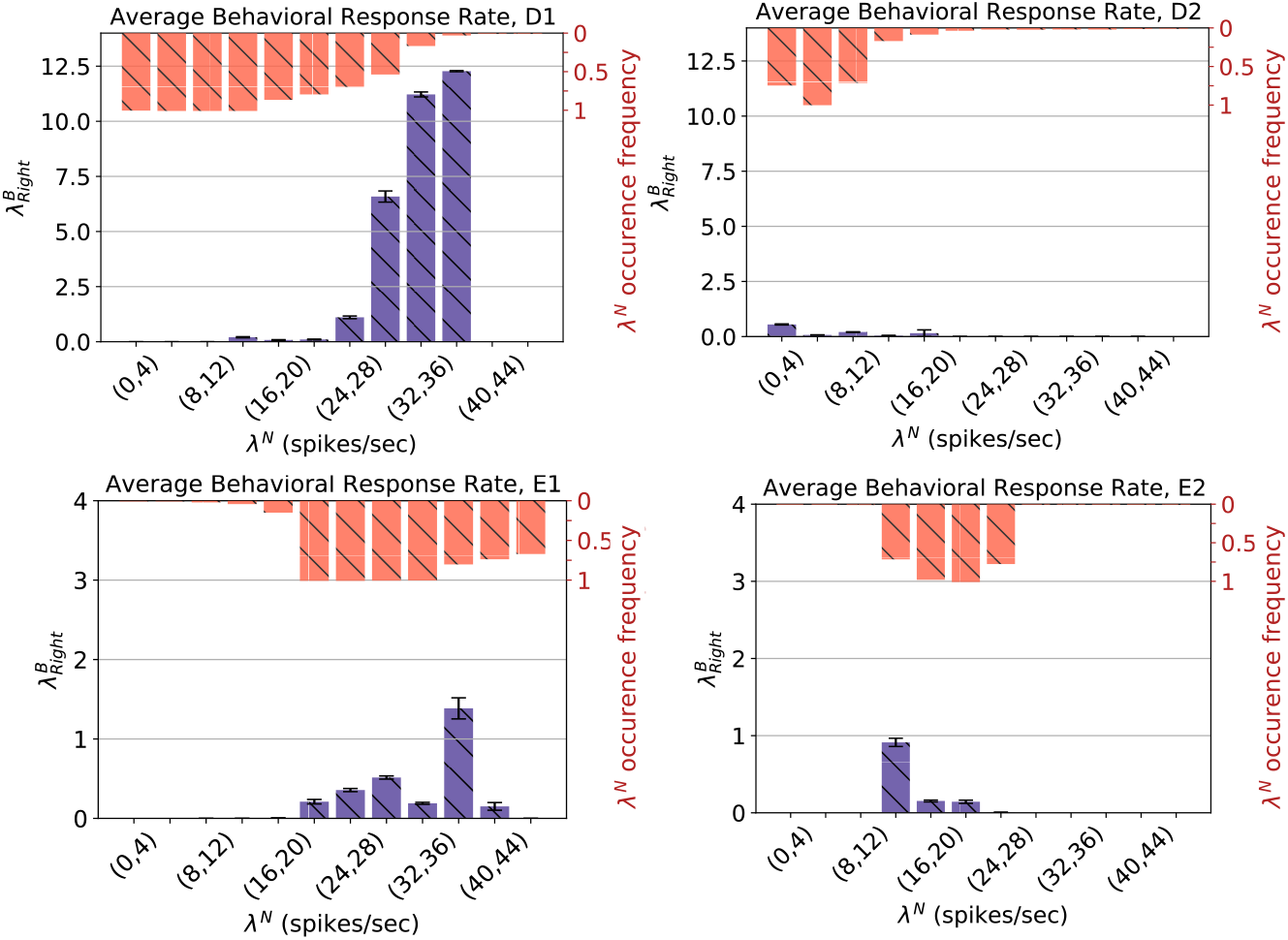
Synthetic Dataset. Illustrating the average estimated response rate for each interval of estimated neural activity. Error bars show the standard error. Purple bars show the average behavioural response rates. Orange bars indicate the proportion of trials with the occurrence of each neural response interval. As expected in the right trials and represented by purple bars, highest behaviour rates are detected in D1 when neural activities approach 40 spikes/s in the (32,36) and (36,40) range. On the other hand, there is little to no activity estimated for D2 which is a leftward preferring integration region. Highest response rates are achieved for when E1 neural activity is in the vicinity of 40 spikes/s (in the (36,40) range) and when the neural activity in E2 is in (12,14) range. These are in agreement with results from Figure 8.

#### G.2 Steinmetz dataset

With a similar procedure as explained above, Figure 10 shows the comparison of the model estimation with the empirically derived firing rates for all the regions in the Steinmetz dataset. The occasional observed underfitting for some regions may be overcome by further forms of regularization in future works. The plots highlight the performance quality of the trained network on test regions. The temporal neural pattern in each region is well captured by the model outputs for different observation windows *W_n_*.

The behavioural performances were also evaluated similar to above for the available test regions in the Steinmetz dataset. Figure 11 illustrates the results. In each region, the highest response rates correspond to the neural activity values achieved near the end of trial in Figure 10. The purple bars show the average behavioural response rate in each neural activity interval. The orange bars show how frequently these were observed in each right trial; meaning the occurrence ratio of that specific neural response range over all right test trials.

**Figure 10:**
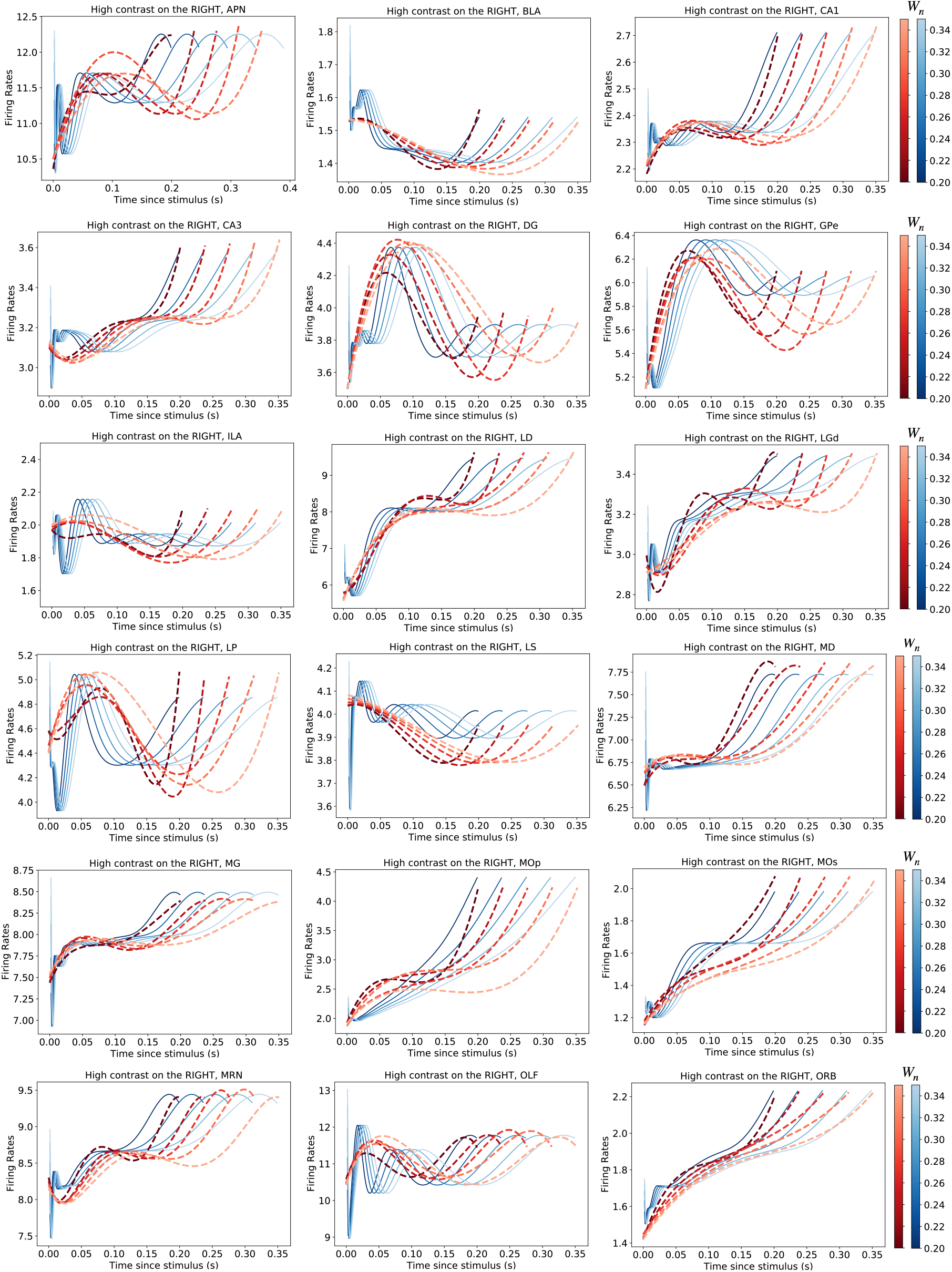
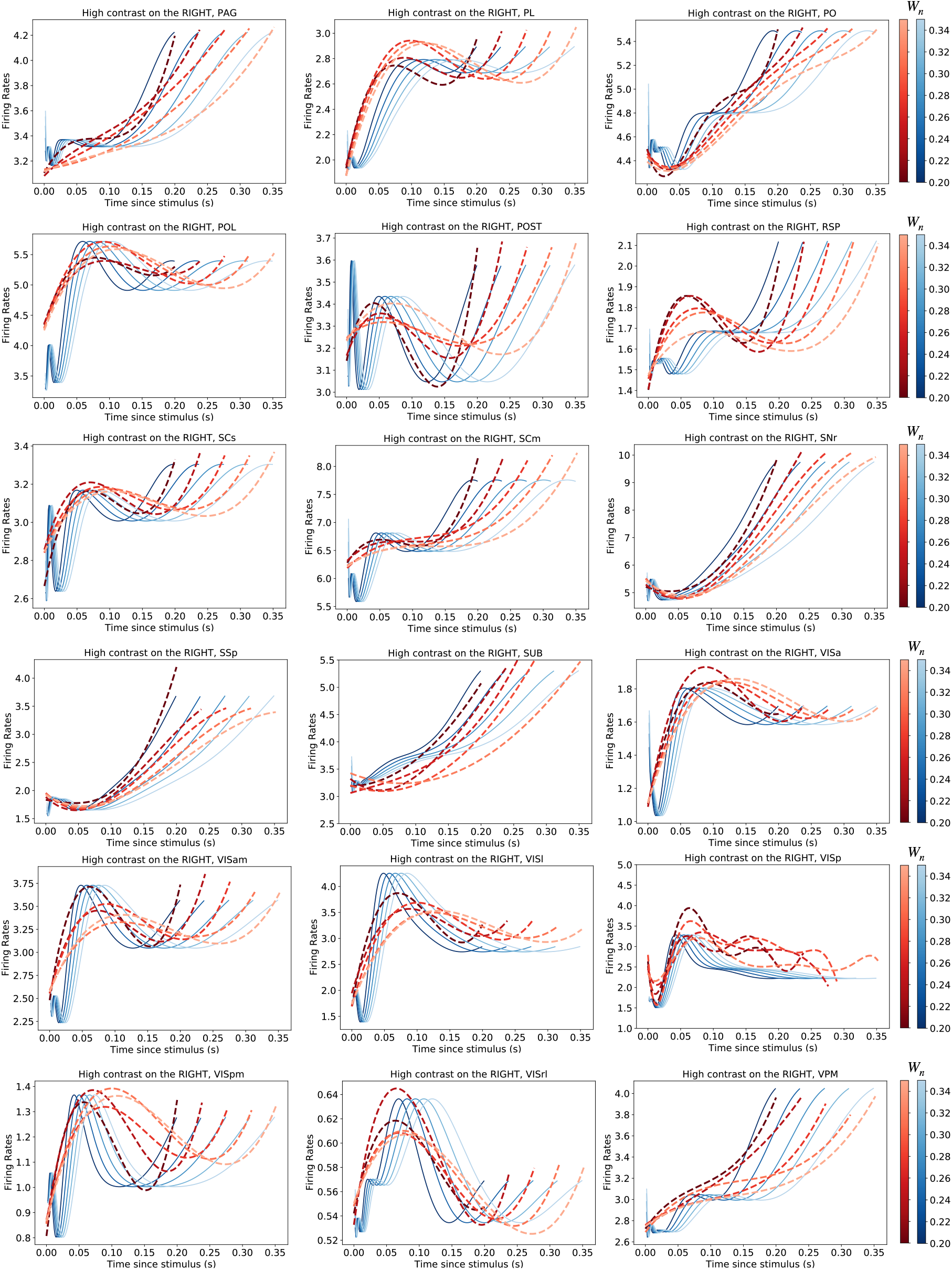
Steinmetz Dataset. The red dashed lines are the empirically derived firing rates compared to the blue solid lines which are estimated using the proposed model. The plots correspond to the trials with high contrast on right. The results are obtained only using trials with the right side contrast higher than left.

**Figure 11:**
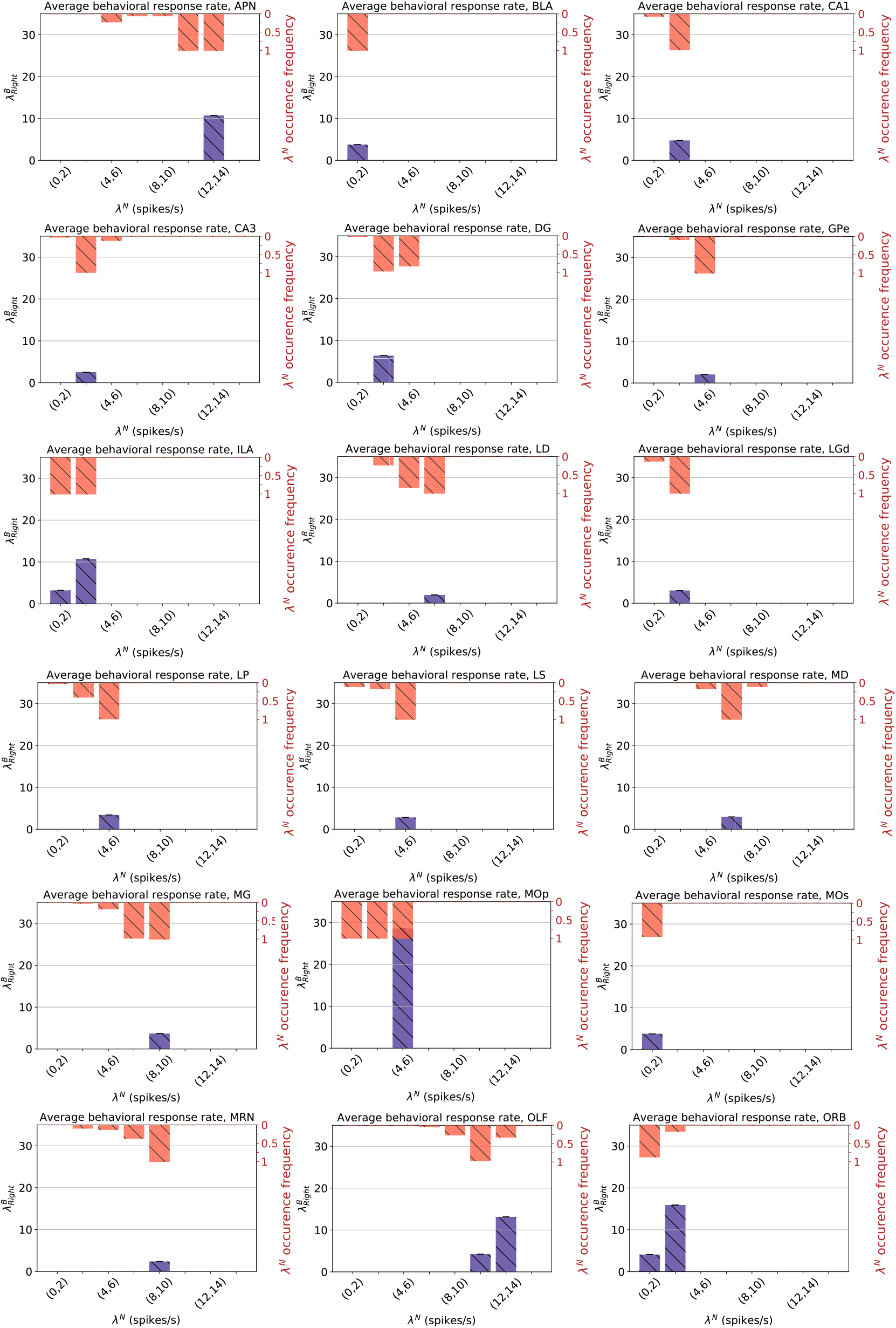
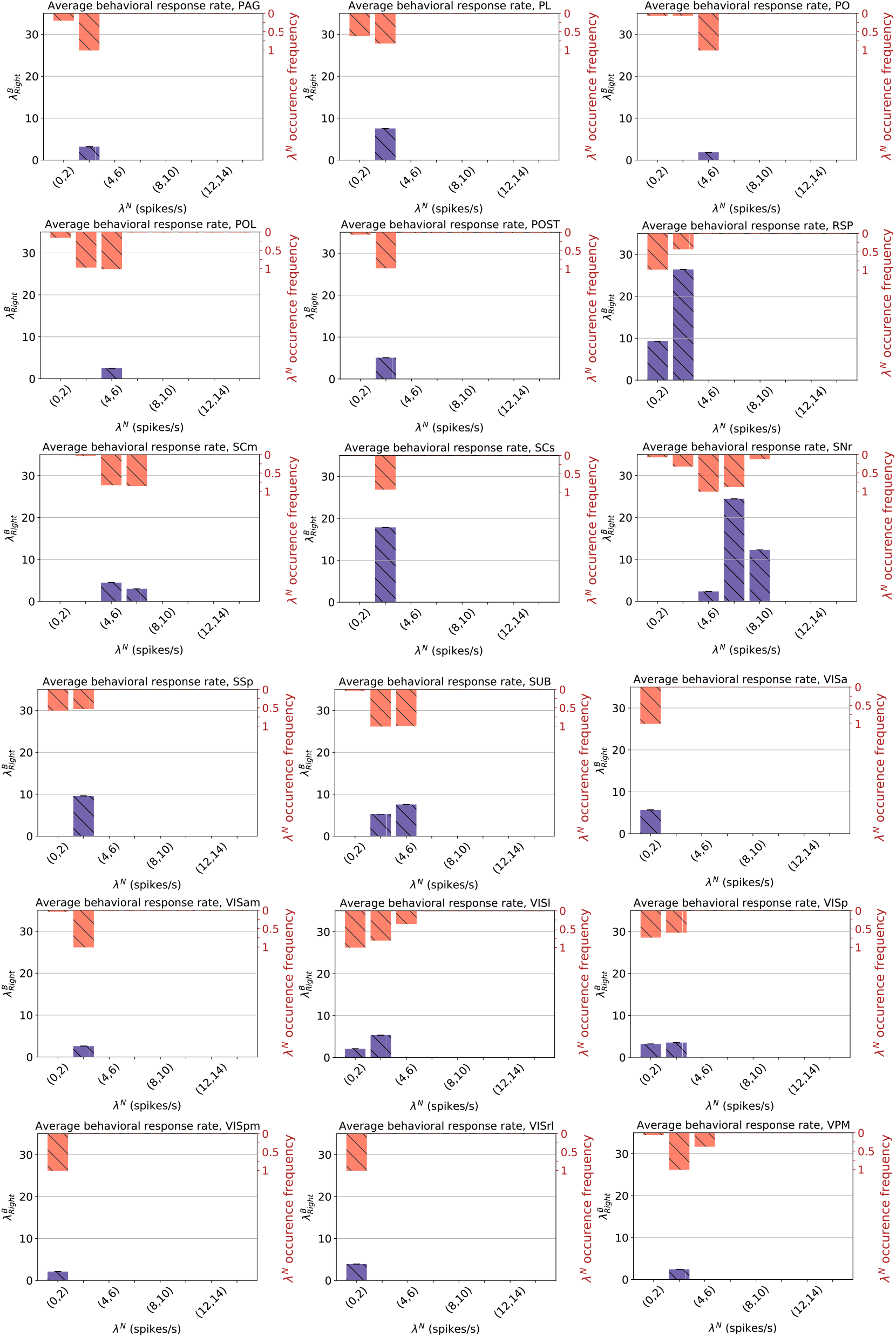
Steinmetz Dataset. Illustrating the average estimated response rate for each interval of estimated neural activity. The purple bars show the average behavioural response rate and orange bars represent the occurrence ratio of that specific neural response interval among all right test trials. Error bars show the standard error. The size of error bars is invisible compared to the bar sizes in most cases.

## Footnotes

2 https://github.com/nsteinme/steinmetz-et-al-2019/wiki/data-files

3 https://senselab.med.yale.edu/MicrocircuitDB/showModel.cshtml?model=168867&file=%2Fhierarchical_network%2Freadme.html#tabs-2

4 https://github.com/nsteinme/steinmetz-et-al-2019/wiki/data-flles

## Notes

### Competing Interest Statement

The authors have declared no competing interest.

